# Age-dependent increased sag amplitude in human pyramidal neurons dampens baseline cortical activity

**DOI:** 10.1101/2021.11.03.467014

**Authors:** Alexandre Guet-McCreight, Homeira Moradi Chameh, Sara Mahallati, Margaret Wishart, Shreejoy J. Tripathy, Taufik A. Valiante, Etay Hay

**Affiliations:** Krembil Centre for Neuroinformatics, Centre for Addiction and Mental Health; Krembil Brain Institute, University Health Network; Department of Psychiatry, University of Toronto; Institute of Medical Sciences, University of Toronto; Department of Electrical and Computer Engineering, University of Toronto; Institute of Biomaterials and Biomedical Engineering, University of Toronto; Department of Surgery, University of Toronto; Center for Advancing Neurotechnological Innovation to Application; Max Planck-University of Toronto Center for Neural Science and Technology; Department of Physiology, University of Toronto

## Abstract

Aging involves various neurobiological changes, although their effect on brain function in humans remains poorly understood. The growing availability of human neuronal and circuit data provides opportunities for uncovering age-dependent changes of brain networks and for constraining models to predict consequences on brain activity. Here we found increased sag voltage amplitude in human middle temporal gyrus layer 5 pyramidal neurons from older subjects, and captured this effect in biophysical models of younger and older pyramidal neurons. We used these models to simulate detailed layer 5 microcircuits and found lower baseline firing in older pyramidal neuron microcircuits, with minimal effect on response. We then validated the predicted reduced baseline firing using extracellular multi-electrode recordings from human brain slices of different ages. Our results thus report changes in human pyramidal neuron input integration properties and provide fundamental insights on the neuronal mechanisms of altered cortical excitability and resting state activity in human aging.

## Introduction

Aging of the human brain is a variable and multifaceted process in terms of cognitive, cellular and anatomical changes (Peters, 2006). Among the different age-associated declines in cognitive performance, older people often exhibit slower cortical processing (Bucur & Madden, 2010) and reduced discrimination acuity (Legge et al., 2008, 2019). Whereas the underlying cellular and circuit mechanisms that cause these deficits remain unknown, studies in monkeys indicate that a decline in working memory with age is associated with reduced baseline and response spike rates in prefrontal cortex, which may involve increased hyperpolarization-activated cyclic nucleotide– gated (HCN) current, or h-current (Ramos et al., 2006; Wang et al., 2007, 2011). The increasing availability of human neuronal transcriptomic and electrophysiological datasets (Gouwens et al., 2018; Hodge et al., 2019; Chameh et al., 2021; Goriounova et al. 2018) offers an opportunity to look for neuronal changes across age demographics and, through computational modeling, determine their impacts on signal processing and behavior.

In cortical pyramidal neurons, h-current modulates signal integration, particularly in the apical dendrites due to increased density with distance from soma (Kole et al., 2006). Increased h-current, which is the primary contributor to generating and shaping sag voltage, results in a more depolarized resting membrane potential (Williams & Stuart, 2000; Beaulieu-Laroche et al., 2018) and decreased input resistance (Song & Moyer, 2017), which consequentially dampens excitatory post-synaptic potential (PSP) summation (Williams & Stuart, 2000; Beaulieu-Laroche et al., 2018). Moreover, increased sag voltage amplitude and depolarized resting membrane potentials have been reported in pyramidal neurons of deeper layers in human cortex (Chameh et al., 2021). Previous studies suggest age-associated changes in h-current which could underlie cognitive decline (Ramos et al., 2006; Wang et al., 2007, 2011), whereby blockade of HCN channels can rescue age-associated declines in both working memory and associated spike rates in prefrontal cortex (Wang et al., 2007). Conversely, in mice, deletion of HCN1 channel subunit expression or h-current block in prefrontal cortex leads to a reduction in the prevalence of persistent spiking and deficits in working memory (Thuault et al., 2013), indicating that h-current also supports cortical function.

There is increasing data of h-current properties in human neurons as measured by transcriptomics and sag voltage recordings. Recent studies showed that HCN channel subunits are more ubiquitously expressed in pyramidal neurons across cortical layers of humans relative to rodents, with larger sag voltage in deeper layers compared to superficial layers (Kalmbach et al., 2018; Chameh et al., 2021; Rich et al., 2021). However, it is unknown if sag changes with age in human neurons as suggested by the above studies in monkeys.

We analyzed sag voltage recordings from human cortical layer 5 pyramidal (L5 Pyr) neurons of younger and older subjects to check for age-associated changes in h-current and membrane properties. We then developed younger and older human pyramidal neuron models using these electrophysiological datasets, which we incorporated into microcircuit models. We simulated baseline and response activity in these human microcircuit models to characterize the effects of age-associated changes in Pyr neuron sag amplitude on cortical processing and resting state activity.

## Methods

### Electrophysiology data

We used whole-cell recordings of human middle temporal gyrus layer 5 (L5) pyr neurons reported previously in datasets from Krembil Brain Institute (KBI; Chameh et al., 2021; Younger: n = 60 neurons; Older: n = 13 neurons; see Table S1 for further breakdown of neuron recording counts across male/female) and Allen Brain Institute (ABI; Gouwens et al., 2018; Younger: n = 50 neurons; Older: n = 14 neurons; see Table S2 for further breakdown of neuron recording counts across male/female). Subject age in the KBI and ABI datasets ranged between 21 - 58 years, and 23 - 83 years, respectively. We grouped the subjects into younger (< 50 years) and older (≥ 50 years) in line with previous studies (Davis et al., 1990; Bucur & Madden, 2010; McIntosh et al., 2014). We also used intracellular recordings of human putative L5 parvalbumin (PV, neuron ID: 528687520) interneurons available from ABI (Gouwens et al., 2018). We used reconstructed human neuron morphologies from ABI for the L5 Pyr neuron (Neuron ID: 562381210; donor age = 34 years, female) and the PV interneuron. The L5 pyramidal neuron morphology with fully-reconstructed dendrites that we chose was a typical intratelencephalic-projecting (IT)-like, which is the most abundant Pyr neuron type in human cortical layer 5 (Chameh et al., 2021; Hodge et al., 2019; Kalmbach et al., 2021). We used another morphology that had fully-reconstructed dendrites (Neuron ID: 529863215; donor age = 67 years, male) to test the generalization of the modeling results. We used depolarizing and hyperpolarizing current steps for neuron model optimization (**Tables S3-S5**), whereby Pyr neuron data was from the KBI dataset because it included a larger range of hyperpolarizing and depolarizing step current amplitudes (i.e., as low as -400 pA and as high as 300 pA). Three depolarizing supra-threshold steps were used for fitting spiking features at low, medium and high firing rates. The hyperpolarizing step currents were used for fitting passive and sag features.

For data collection for the KBI dataset (Chameh et al., 2021), we visualized the cortical layers using an IR-CCD camera (IR-1000, MTI, USA) with a ×40 water immersion objective lens. Using the IR-DIC microscope, the boundary between layer 1 and 2 was easily distinguishable in terms of cell density. Below L2, the sparse area of neurons (L3) was followed by a tight band of densely packed layer 4 (L4) neurons. L4 was followed by a decrease in cell density (L5). Following electrophysiological recording, we confirmed human cortical layers using DAPI-staining on sections (500 µm) cleared with the CLARITY technique.

We also used extracellular recordings using MEA (electrode pitch: 200-300 micrometer; TiN electrode array – Multi Channel 151 Systems, Germany) from resected human middle temporal gyrus slices (500 μm thickness) maintained active in carbogenated (95% O_2_, 5% CO_2_) artificial cerebral spinal fluid (Florez et al., 2015; Chameh et al., 2021). The electrodes were registered on the slices to identify the anatomical location of the single units recorded. Putative cell types were identified based on waveform features, mainly the peak to trough latency of the average spike waveform (Barthó et al., 2004). We used the waveform feature to cluster units into broad spiking and narrow spiking units. Subsequent spike rate analyses were performed on broad-spiking units (putative pyramidal neurons). Subject age ranged between 19 – 65 years and were grouped subjects into younger (< 50 years) and older (≥ 50 years) groups (Younger: n = 376 L2/3, 119 L4, and 401 L5 units from 17 subjects; Older: n = 148 L2/3, 37 L4, and 51 L5 units from 5 subjects). While we had L6 data, we did not include this age comparison because of too little data in the older group.

### Electrophysiological feature analysis

To maximize our usage of the datasets, we z-scored the raw sag amplitudes respective to the means and standard deviations at each current step (where +/-z-score values indicated sag amplitudes above or below the mean of the corresponding current step) and then pooled the data across current steps accordingly. The data was then grouped by demographic parameters (i.e. age and/or gender). This method was supported by the z-scored metrics exhibiting a flat relationship with current magnitude and across current steps. We performed similar z-scoring for other features such as sag ratio and input resistance. All the above features were computed using functions from the Electrophys Feature Extraction Library (eFEL), and sag ratio was computed as the ratio of sag amplitude and the maximal deflection.

### Statistical tests

For data that exhibited significantly non-uniform distributions (Omnibus test of normality, *p* < 0.05) and unequal variances between younger and older groups (Levene test, *p* < 0.05), we used Welch’s t-tests for between group comparisons (Ruxton, 2006; Skovlund & Fenstad, 2001). Otherwise, we used paired-sample t-tests or Mann Whitney U tests where applicable. To estimate how age and other covariates were associated with sag amplitude, we ran analyses of linear mixed-effect models in R using the lmer function from the lmerTest package (Kuznetsova et al., 2017) with restricted maximum likelihood, where subject and cell identifiers were modelled as random effects and either age or age group were modelled as fixed effects (*Sag amplitude* ∼ *Age* + (1| *Subject ID*) + (1|*cell ID*)). We conducted similar tests to account for multiple spiking units from the same subject (*Spike rate* ∼ *Age* + (1| *Subject ID*)). We then used likelihood ratio tests to compare the full models against the corresponding reduced models with the fixed effects dropped.

### Human L5 microcircuit models

We simulated L5 microcircuits comprised of 1000 neurons distributed along a layer 5 volume (500×500×700 μm^3^, 1600 to 2300 μm below pia; Mohan et al., 2015) using NEURON (Carnevale & Hines, 2006) and LFPy (Hagen et al., 2018). In addition to Pyr and PV neurons, the microcircuit models included somatostatin (SST) and vasoactive intestinal peptide (VIP) interneurons, using previous human L2/3 neuron models (Yao et al., 2022), because there was no human data for these interneurons from L5. Neurons of a given type had the same model (morphology and biophysical properties) but differed in the randomization of their synaptic connectivity and background input. The proportions of the four neuron types in the microcircuit were: 70% Pyr, 15% SST, 10% PV, and 5% VIP. These were approximated using ultra high-depth human neocortex single-nucleus RNA-seq data from the Allen Institute for Brain Sciences “Multiple Cortical Areas - Smart-seq (2019)” dataset (https://portal.brain-map.org/atlases-and-data/RNA-seq/human-multiple-cortical-areas-smart-se), with sample collection and data analysis methodologies described previously (Hodge et al., 2019). Specifically, we used data from the upper and lower limb primary somatosensory cortical regions. In general, for microcircuit model parameters we used human cellular and circuit data when available, and data from rodents or monkeys when human data was not available (summarized in **Table S6**).

### Human neuron models

We developed multi-compartmental models for the Pyr and PV neurons using the BluePyOpt Python module to perform multi-objective optimizations (Van Geit et al., 2016). We used a set of ion channel mechanisms taken from previously published models (Hay et al., 2011, 2013; Yao et al., 2022). The models were fit in one step where all passive and active parameters and features were optimized simultaneously (**Tables S3-S5**). We set the following parameters: axial resistance (*R*_*a*_) = 100 Ω cm, sodium reversal potential (*E*_*Na*_) = 50 mV, potassium reversal potential (*E*_*K*_) = *-85* mV, and percent of free calcium (*CaDynamics*_*gamma*_) = 0.0005 (Hay et al., 2011). We also set the somatic and axonal Na_T_ kinetics controlling the half voltage (*Vshift*) and slopes of the voltage steady-state activation (*m*) and inactivation (*h*) sigmoidal functions to *Vshift*_*m*_ = 0, *Vshift*_*h*_ = 10, *Slope*_*m*_ = 9, and *Slope*_*h*_ = 6. The specific membrane capacitance (*c*_*m*_) was 0.9 µF/cm^2^ for the Pyr models, and 2 µF/cm^2^ for the PV model to reproduce membrane time constants (possibly due to errors in PV dendritic diameter estimation; **Table S7**). Model optimization was run using parallel computing clusters [Neuroscience Gateway (NSG; Sivagnanam et al., 2013) & Scinet (Loken et al., 2010; Ponce et al., 2019): 400 processors with a population size of 400, across 300 generations and an approximate total runtime of 5 hours]. Ion channels were inserted primarily in the soma and axon initial segment compartments, where we included some of the same somatic objective experimental targets to constrain the axonal spiking (**Tables S3-S5**). Additionally, 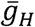 was distributed uniformly across all dendritic sections of the PV model (similarly to the SST and VIP models). For Pyr neurons, we have adapted an exponential function used in previous models (Hay et al., 2011) to an equivalent sigmoidal function allowing saturation at a certain distance from soma, so that 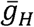 increased with distance from soma along the basal and apical dendrites as follows:

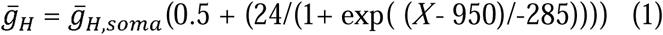

where *X* is the distance from soma in μm. Model performance in reproducing the electrophysiology features was assessed in terms of standard deviation from the experimental mean. For Pyr models we used the statistics over the set of recorded neurons from the KBI dataset, and for the PV model (where data from only a single human neuron was available), we used the population variance from the rodent literature (Zurita et al., 2018).

### Synaptic connectivity models

We used previous models of NMDA/AMPA excitatory and GABA_A_ inhibitory synapses, that incorporated presynaptic short-term plasticity parameters for vesicle-usage, facilitation, and depression, as well as separate time constant parameters for the AMPA and NMDA -mediated components of excitatory synapses (Fuhrmann et al., 2002; Hay et al., 2013; Mäki-Marttunen et al., 2019). We set the same time constant parameters for all connection types (τ_*rise,NMDA*_ = 2 ms; τ_*decay,NMDA*_ = 65 ms; τ_*rise,AMPA*_ = 0.3 ms; τ_*decay,AMPA*_ = 3 ms; τ_*rise,GABA*_ = 1 ms; τ_*decay,GABA*_ = 10 ms), as well as the reversal potential values (*E*_*exc*_ = 0 mV; *E*_*inh*_ = - 80 mV).

For Pyr→Pyr connections, we fitted the synaptic conductance and vesicle-usage parameters using the human experimental literature (Seeman et al., 2018). We simulated the experimental conditions (e.g., chloride reversal potential, and holding currents) and adjusted the conductance and vesicle-usage parameters to achieve the target postsynaptic (PSP) amplitudes and failure rates on average across 50 randomizations of synaptic locations and events. For the conductance and vesicle-usage parameters of all other connection types, as well as the depression, facilitation, and numbers of synaptic contacts (*N*_*syns*_) for all connections, we used values reported by the Blue Brain Project (Ramaswamy et al., 2015). For SST→Pyr connections, we used a higher conductance, consistent with SST→Pyr conductance of human synaptic connections in L2/3 (Yao et al., 2022). The synaptic parameters of the different connections are summarized in **Table S8**. Specific synaptic locations onto Pyr neurons were dependent on the connection type, where Pyr→Pyr synapses were placed on both basal and apical dendritic compartments, PV→Pyr connections were placed on basal dendritic compartments, and SST→Pyr connections were placed on apical dendritic compartments. Apart from these specifications, synapse locations were chosen randomly from a uniform distribution.

We initially set unidirectional connection probability (*p*_*con*_) according to the rodent literature (Blue Brain Project and ABI), except for Pyr→Pyr connections, where *p*_*con*_ could range from 7% to 12% according to human literature (Campagnola et al., 2022; Seeman et al., 2018). We adjusted connection probabilities guided by the reported experimental ranges to reproduce the intrinsic activity, which required a Pyr→Pyr connection probability of 9%. The connection probability for the different types of connections in the microcircuit, along with the synaptic conductance (*G*_*syn*_), number of contacts per connection, relaxation time constants from facilitation, relaxation time constants from depression, and utilization of synaptic efficacy (*use*) parameters, are all summarized in **Table S8**.

### Modeling microcircuit intrinsic activity

We constrained the microcircuit to generate baseline spike rates within range and close to the medians recorded for different L5 neuron types in rodents *in vivo* (range, median; Pyr: 0.4 - 11.5 Hz, 2.6 Hz; PV: 4.6 - 22.0 Hz, 17.2 Hz; SST: 0.5 - 6.9 Hz, 1.7 Hz; VIP: 8.5 – 21.0 Hz, 11.1 Hz; Yu et al., 2019), by adjusting the *p*_*con*_ values guided by the reported experimental ranges for all connection types, and by adjusting the background input (see below). We modeled L5 Pyr neurons as a homogenous population, and did not differentiate between subtypes (intratelencephalic-projecting type 1 and type 2, and extratelencephalic-projecting), and the experimental target rates did not differentiate subtypes either. We calculated the simulated rates across non-silent neurons (> 0.2 Hz) over 4.5 seconds of baseline simulation. Average rates were then computed across 30 randomized microcircuits. The microcircuit received random uncorrelated background excitatory input using Ornstein-Uhlenbeck (OU) point processes (Destexhe et al., 2001), placed at halfway the length of each dendritic arbor to ensure similar levels of inputs along each dendritic path. For the Pyr models, we placed 5 additional OU processes along the apical trunk at 10%, 30%, 50%, 70%, and 90% of the apical dendritic length. We set the base excitatory OU conductance to the following: Pyr = 76 pS; SST = 32 pS; PV = 1545 pS (due to higher rheobase); VIP = 75 pS. We did not use an inhibitory OU conductance since the model microcircuit provided sufficient inhibition. Furthermore, we scaled the OU conductance values to increase with distance from soma by the multiplying them with the exponent of the relative distance from soma (ranging from 0 to 1): 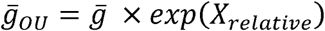.

### Tonic inhibition

We modelled tonic inhibition using a previous model for outward rectifying tonic inhibition (Bryson et al., 2020). We used previous estimates of tonic inhibition conductance (*G*_*tonic*_; uniformly across all somatic, basal, and apical compartments) from a human L2/3 microcircuit model (*G*_*tonic*_: 0.938 mS/cm^2^ for all neuron types; Yao et al., 2022), since total tonic inhibition current in L2/3 Pyr neurons was similar to estimates of the total tonic inhibitory current measured in L5/6 neurons (Scimemi et al., 2006). As well, it has previously been shown that Pyr neurons and interneurons in L5/6 have similar levels of tonic inhibition after correcting for cell capacitance (Scimemi et al., 2006).

### Modeling microcircuit response activity

We modelled Pyr neuron responses according to previous studies of somatosensory cortex in awake monkeys during tactile stimulation of the finger by indentation with an edge at varying orientations (Thakur et al., 2006; Bensmaia et al., 2008). We constrained neuronal tuning curve maximal rates in the range of 10 to 120 Hz (Thakur et al., 2006; Bensmaia et al., 2008), and tuning curve half-width in the range of 34° to 49° (Bensmaia et al., 2008). We applied orientation-selectivity to 50% of the Pyr neurons (Bensmaia et al., 2008), as well as the PV and VIP neurons (Sermet et al., 2019). Pyr basal dendrites, PV neurons, and VIP neurons were stimulated using excitatory AMPA/NMDA synapses with the same synaptic dynamics and conductance as the intra-cortical excitatory synapses above. We injected the thalamic input synapses with noisy artificial presynaptic inputs spiking at 100 Hz, with inter-spike intervals randomly sampled from a negative exponential distribution, over the course of a 100 ms window, in line with the spiking of ventral posterior nucleus neurons (which provide inputs to somatosensory cortex) recorded in awake monkeys during finger indentations (Song & Semework, 2015). Average response rates were calculated over the 100 ms window following stimulus onset. We did not tightly constrain the response spike rates of the interneurons due to a lack of experimental data, but rather constrained the PV and VIP neurons to be activated, while SST neuron response rates were either unchanged or silenced (Gentet et al., 2012; Yu et al., 2019; Sermet et al., 2019; Muñoz et al., 2017).

Neurons in the microcircuit had different preferred orientations, so that angles from 0 - 180° were represented uniformly across Pyr neuron number 1 – 350. The tuning curve of each neuron followed a Gaussian distribution with a standard deviation of 42° (Bensmaia et al., 2008), and was implemented by varying the number of thalamic input synapses (at peak of gaussian: N_syn,Pyr_ = 45 synapses). A similar stimulation paradigm was applied also to the PV and VIP interneurons (peak N_syn,PV_ = 30 synapses, N_syn,VIP_ = 30 synapses), although with a broader tuning curve (standard deviation of 147° and 294°, respectively; Wang et al., 2004) which allowed sufficient activation of PV and VIP interneurons for SST neuron suppression (Gentet et al., 2012; Muñoz et al., 2017). Across random seeds, we additionally applied a non-systematic noise of ±30% (sampled from a uniform distribution) to the number of thalamic input synapses, corresponding to input errors e.g. due to finger position and in tactile surface irregularities, as well as noise in the signal progression along the CNS hierarchy from the periphery. We also tested a condition where synapses were placed on apical dendrites instead of basal dendrites, using 67 synapses to elicit an equivalent average response rate as in the regular basal input condition of the microcircuits with younger Pyr neurons. In addition, we tested a condition of low basal or apical input, using 31 and 45 synapses, respectively.

### Classifier models

Discrimination accuracy using the microcircuit model outputs to 85° vs. 95° stimulus orientations was assessed for microcircuits with younger vs older Pyr neuron models using classifier models available in the scikit-learn Python module. We used angles symmetrically close to 90° because this angle corresponded to the middle-preference for the neuronal population. For each microcircuit model with younger and older Pyr neurons, stimulus discrimination was tested for a single circuit (i.e., fixed connectivity between the neurons) across 80 randomizations of background OU excitation and stimulus amplitude (*N*_*syn*_). Output PSPs from each Pyr neuron during response (0 - 100 ms post-stimulus) were first computed by convolving the neuron output spike trains with an averaged AMPA (τ_*r*_ = 0.5 ms; τ_*d*_ = 3 ms) and NMDA (τ_*r*_ = 2 ms; τ_*d*_ = 65 ms) PSP waveform:

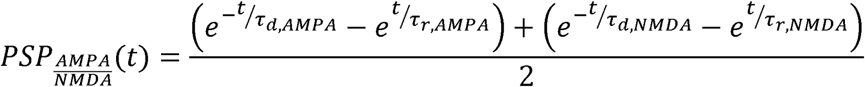

We then computed the area under the curve of the resulting PSPs for each Pyr neuron and used it as feature input to the classifier models. We trained and tested with three different types of classifier models: 1) a linear support vector model, 2) a Gaussian naïve Bayes model, and 3) a multi-layer perceptron model (see **Table S9** for full descriptions of classifier model input arguments). Training was done using a random subset of 15 of the 80 stimulus presentations per orientation and testing was done on the other 65. We derived performance statistics for the classifiers using 300 random permutations of the stimulus presentations used for training and testing. We chose PSP area under curve as the feature inputs because this measure is representative of the amount of charge that downstream postsynaptic neurons will receive.

### PSP summation simulations

To investigate the effects of h-channel density changes with age on synaptic integration, we simulated PSP summation in the younger and older neuron models using a 50 Hz or 25 Hz train of 5 step current pulses (2 ms, 3 nA each) injected at a presynaptic neuron soma (Pyr, SST, or PV) to trigger a train of action potentials and consequently PSPs in a Pyr neuron. Connectivity parameters to the postsynaptic Pyr neuron were the same as in circuit simulations (**Table S8**), except for Pyr→Pyr neuron conductance, which was reduced to 0.3 nS to prevent the postsynaptic Pyr neuron from spiking. We simulated 20 trials (with randomized synaptic probability) across 20 connections (randomized synaptic locations) and calculated the mean and 95% confidence intervals across connections by bootstrapping (500 iterations) the trial means. We ran these simulations with and without DC current compensating for resting membrane potential differences between the younger and older neuron models. We compensated the resting membrane potential by injecting positive current in the younger Pyr neuron (6.65 pA) and negative current in the older Pyr neuron (−6.65 pA). We used two metrics for assessing PSP summation – peak PSP amplitude and area under the PSPs.

### Human HCN1 channel transcriptomics data

We accessed transcriptomic data from the MTG of 8 human postmortem brains (Hodge et al., 2019; https://portal.brain-map.org/atlases-and-data/rnaseq/human-mtg-smart-seq; younger: n = 5 donors, ages 24 – 48 years; older: n = 3 donors, ages 50 – 66 years). Data frames were prefiltered in Python to only include gene counts from glutamatergic neurons located in L5 and to combine intron and exon gene count data frames. We analyzed gene expression using the Seurat v3 toolkit in R (Stuart et al., 2019), where we log-normalized gene counts by dividing each count by the total count for that cell, scaling this by a factor of 10,000, and then computing the natural log-transform using the log1p function. Log-normalized counts for HCN1 transcripts were compared between younger and older cells using a Wilcoxon Ranked Sum test, and the log-fold change of average expression was computed.

### Model population analysis

We analyzed the population parameter sets of younger and older Pyr neuron models generated from our multi-objective optimizations by first selecting sets of acceptable models, across the entire optimization history, based on performance (e.g., within 2 SD from the target experimental means, except 3 SD for AP width, AHP depth features, and AHP slow time). To narrow down our selection to models that captured both sag voltage amplitude and mean frequency well, we set the SD thresholds for these features to less than 1 SD. Models were then sorted by overall error scores and duplicate models were removed. After selecting these sets of models, we normalized their conductance parameter values by the upper and lower search limits used in the optimization (0 corresponding to the lower limit and 1 corresponding to the upper limit).

### Microcircuits with heterogenous Pyr neuronal models

We tested the robustness of our results using microcircuit simulations that had heterogenous models for Pyr neurons, sampled randomly from the top 30 highly-ranked models among the model population sets (see above).

## Results

### Increased sag amplitude in older L5 human pyramidal neurons

We first analyzed sag amplitude in human L5 Pyr neurons (**Fig. 1A**) from the Krembil Brain Institute (KBI) and Allen Brain Institute (ABI) datasets across subject age, and in younger (< 50 years) vs older (≥ 50 years) groups. The sag amplitude in the KBI dataset increased with age (Pearson correlation, R = 0.31, *p* < 0.05, **Fig. 1B**), and also between younger and older subject groups (two-sample Welch’s t-test, *p* < 0.05, Cohen’s *d* = 1.00; **Fig. 1C**). The increase was seen for example steps (**Fig. 1B-C**) and also when comparing sag amplitudes across all steps (see Methods, two-sample Welch’s t-test, *p* < 0.05, Cohen’s *d* = 0.91; **Fig. 1D**). Sag amplitude across steps increased similarly between younger and older subject groups in the ABI dataset (two-sample Welch’s t-test, *p* < 0.05, Cohen’s *d* = 0.41; **Fig. 1E**). Similar results were seen when comparing sag ratio across age groups.

**Figure 1.**
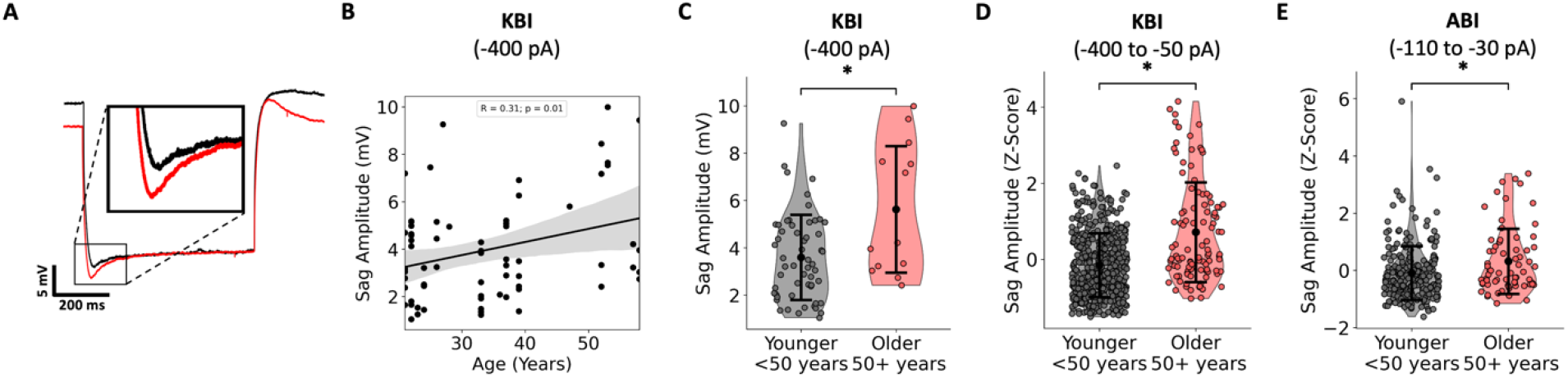
Increased sag amplitude in older human L5 Pyr neurons. **A**. Example experimental voltage response to hyperpolarizing current injection (−400 pA/600 ms) recorded in L5 Pyr neurons from younger (< 50 years, black) and older (≥ 50 years, red) subjects from the KBI dataset. Traces were aligned by steady-state voltage. Inset shows sag voltage comparison. **B**. Sag amplitudes in the KBI dataset increased with age (for example current steps of -400 pA). Shaded area in all regression plots denotes the bootstrapped 95% confidence intervals. **C**. Sag amplitudes were larger in older compared to younger subjects (for example current steps of -400 pA). **D-E**. Sag amplitude increased across all hyperpolarizing current steps (z-scored) in older compared to younger subjects in the KBI dataset (D, -400 pA to -50 pA step amplitudes) and in the ABI dataset (E, -110 pA to -30 pA step amplitudes). Error bars in all plots denote standard deviation.

Variable resting potentials across neurons were not controlled for in either set of experiments but correlated weakly with sag amplitude and ratio. Importantly, none of the additional features we analyzed (i.e., resting potential, input resistance, steady-state voltage, membrane time constant) exhibited consistent changes with age across both datasets. While the effects were strong on the neuronal level, when averaged per subject we only found trends towards increasing sag with age in both datasets (**Fig. S1A**), potentially due to the small subject sample size. However, linear mixed-effect models using age group as fixed effects further indicated that the effects on sag amplitude were driven by age and not cell/subject variability (likelihood ratio test, *p* < 0.05). Similar support was seen in linear mixed-effect models using age as fixed effect (likelihood ratio test, *p* = 0.06).

The increase in sag amplitude in older participants was also seen in gender-specific comparisons, which we were able to conduct using the KBI dataset due to an even distribution of female participants in the younger and older groups and an even distribution of male participants in the younger group (**Fig. S1B**). Sag amplitude in older females was significantly larger than in younger females (two-sample Welch’s t-test, *p* < 0.05, Cohen’s *d* = 0.96, **Fig. S1C**). Although a similar comparison was not possible for male participants, in the younger group males and females had similar sag amplitudes (**Fig. S1D**), further indicating that effects were due to age differences and not gender.

We generated models for younger and older L5 Pyr neuron models using multi-objective evolutionary algorithm optimization, constrained with recordings from the KBI dataset and a human neuron morphology (**Fig. 2**). Younger and older neuron models (n = 57 and 52, respectively) reproduced the experimental spiking features and hyperpolarization sag features (**Fig. 2B-E**; **Tables S3-S4**), whereby all features were within the experimental range (1 - 3 standard deviations, SD, **Tables S3-S4** and **Figs. S2-S3**, see Methods). The only exception was spike width, which was narrower than the experimental range, a typical limitation of the particular ion channel kinetics we have used. Similar quality fits were obtained when optimizing using an alternate reconstructed neuron morphology reconstruction (see Methods, **Fig. S4A-C**). We chose exemplar models for younger and older neurons (**Table S7**) that best reproduced the different features (**Fig. S2, S3**) and had a similar fit quality between younger and older models (**Fig. S4B**). As seen experimentally, spiking features were similar between the younger and older neuron models (**Fig. 2B-C**), but the older neuron model had a larger sag amplitude (**Fig. 2D-E**). Though the older neuron model had a more depolarized resting membrane potential (−69.8 vs. - 71.2 mV, respectively, in line with the experimental population mean targets shown in **Tables S3-S4**), we note that there were no significant differences in resting membrane potential between younger and older pyramidal neuron datasets. Accordingly, the older model neuron had a larger h-channel density, and this difference was seen also when comparing *G*_*H*_ in the set of acceptable models for younger and older neurons derived from the optimization algorithm (**Fig. S4C, top**), as well as when using an alternate morphology (**Fig. S4C, bottom**; Bonferroni adjusted *p* < 0.01, Wilcoxon Ranked Sum test). Due to the increased h-current density along the apical dendrites, the difference between younger and older model neurons was more pronounced in the distal apical dendrites (**Fig. S4D**). This difference in h-channel density was further supported by an increased HCN1 subunit expression in older human MTG L5 excitatory neurons (Benjamini-Hochberg adjusted *p* < 0.01, Wilcoxon Ranked Sum test, Cohen’s *d*: 0.18, log-fold increase: 0.11; **Fig. S4E**).

**Figure 2.**
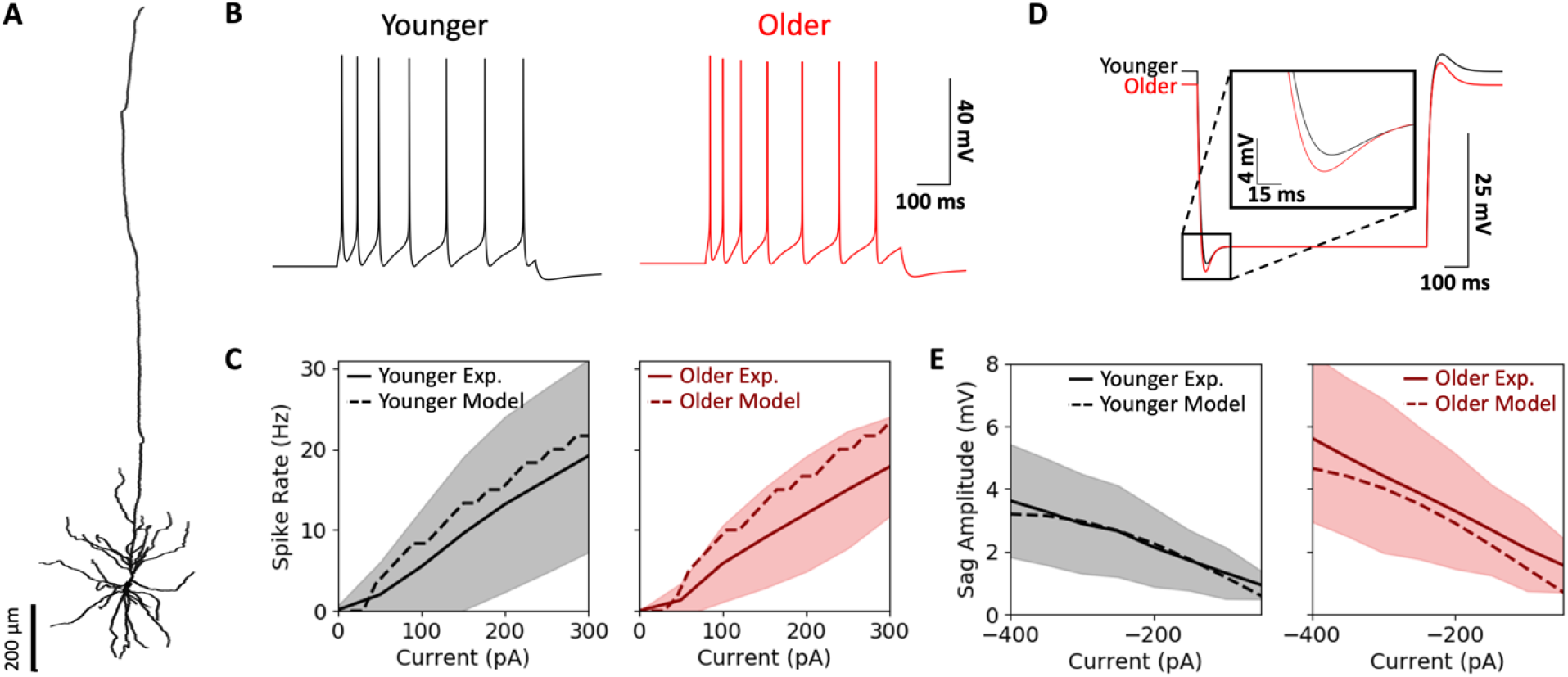
Human L5 Pyr neuron models capture electrical properties in younger and older data. **A**. Reconstructed human L5 Pyr neuron morphology, used in the younger and older neuron models. **B**. Spiking output of younger and older Pyr neuron models in response to a depolarizing step current of 135 pA and duration of 600 ms. **C**. Spike frequency and input current relationship for younger and older models (dashed) was within the experimental range, < 1 SD (shaded) from the experimental mean (solid). **D**. Voltage response to hyperpolarizing step current of -400 pA and duration of 1000 ms in the younger and older models. Traces are aligned according to their steady-state voltages to highlight differences in sag amplitude (inset). **E**. Sag amplitude and input current relationship for younger and older models (dashed) were within the experimental range, < 1 SD (shaded) from the experimental mean (solid).

### Dampened baseline spiking in L5 microcircuit models with older Pyr neurons

We next integrated our younger and older Pyr neuron models into cortical microcircuit models, complete with similarly detailed models of human PV, SST, and VIP inhibitory neurons (**Fig. 3A,B**). We used previous models for human SST and VIP interneurons and fitted models for the human L5 PV interneuron (**Tables S7**). The passive and active firing features of the PV interneuron model were within the experimental variance of population data from corresponding neurons in rodents, except for AP half-width which is more dependent on the particular parameters used for the channel kinetics (Yao et al., 2022). We constrained Pyr→Pyr connections to reproduce experimental EPSP amplitudes reported in human L5 Pyr neurons (model: 0.81 ± 0.69 mV, experimental: 0.80 ± 0.69 mV; Seeman et al., 2018). The remaining connections were constrained using rodent data (**Table S8**).

**Figure 3.**
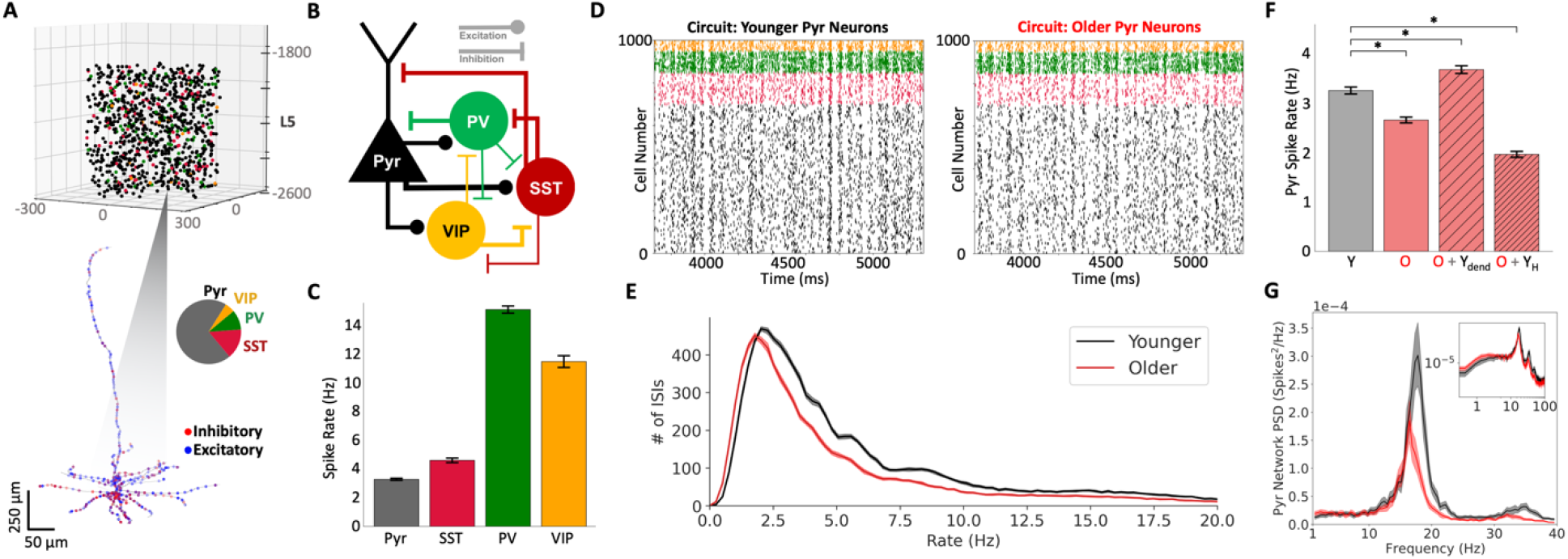
Dampened baseline spiking in human cortical microcircuit models with older Pyr neurons. **A**. The model microcircuit comprised of 1000 neurons, with the somas distributed in a 500×500×700 μm^3^ volume along L5 (1600 to 2300 μm below pia). The proportions of the different neuron types were according to the experimental transcriptomics data in human somatosensory cortical L5 (pie chart, Pyr: 70%; SST: 15%; PV: 10%; VIP: 5%). The neurons of all four types were modeled with detailed morphologies, as in Figure 2. The blue and red dots denote example excitatory and inhibitory synapses, respectively. **B**. Schematic diagram of the key connections between different neuron types in the microcircuit. **C**. Spike rates in the different neuron types reproduced experimental baseline firing rates (mean and standard deviation, n = 30 younger Pyr neuron microcircuits). **D**. Example raster plot of baseline spiking in the microcircuit models with younger and older Pyr neurons, color-coded according to neuron type. **E**. Distributions of instantaneous firing rates of Pyr neurons from younger and older microcircuit models (bootstrapped mean and 95% confidence intervals of 30 microcircuits). **F**. Mean Pyr neuron spike rate (excluding silent Pyr neurons firing at <0.2 Hz) significantly decreased in microcircuit models with older (red; O) compared to younger (black; Y) Pyr neurons (n = 10 microcircuits). Spike rates were recovered when changing the dendritic h-channel density and leak current parameters in the older models to that of the younger models (O + Y_dend_), but not when changing dendritic h-channel density alone (O + Y_H_). **G**. Spikes PSD of Pyr neurons for both microcircuit models (younger, black; older, red), bootstrapped mean and 95% confidence intervals (n = 30 randomized microcircuits). Inset – PSD in log scale, illustrating the 1/f relationship between power and log frequency.

Using the younger Pyr neuron model, we simulated human cortical L5 microcircuits of 1000 neurons with experimentally derived cortical dimensions and proportions of different neuron types (**Fig. 3A, B**). Each neuron received random background excitation corresponding to cortical and thalamic drive, to enable recurrent activity. We tuned the connection probabilities between neuron types and the background excitation levels to reproduce baseline firing rates previously reported for the neuron types *in vivo* (**Fig. 3C, D**). The mean firing rate for simulated Pyr neurons was 3.27 ± 0.07 Hz, PV: 15.03 ± 0.29 Hz, SST: 4.56 ± 0.18 Hz and VIP: 11.36 ± 0.32 Hz (n = 30 randomized microcircuits).

We then simulated L5 microcircuit models using the older Pyr neuron model, and found a downward shift in the distribution of instantaneous baseline Pyr neuron spike rates (**Fig. 3E**, Mann-Whitney U, *p* < 0.001) and a decrease in mean Pyr neuron spike rates (**Fig. 3F**, paired-sample t-test, *p* < 0.001, Cohen’s *d*: -8.7). We also tested this effect when changing the h-channel mechanism kinetics to values that have been used in previous modeling work (Hay & Segev, 2015; Rich et al., 2021), and consistently obtained decreased baseline Pyr neuron spike rates. This age effect on microcircuit firing remained consistent also when simulating microcircuits with heterogeneous Pyr neuronal models (see Methods) or a larger network size of 2600 neurons (paired-sample t-tests, *p* < 0.01; Cohen’s *d* = -13.8 and -3.5, respectively). This decrease in baseline spike rate could be recovered and even increased by changing dendritic h-channel density and passive membrane parameter values in the older Pyr neuron model dendrites to the values used in the younger Pyr neuron model dendrites (**Fig. 3F**, paired-sample t-test, *p* < 0.001; Cohen’s *d*: 5.4), indicating that the change in spike rate was primarily driven by differences in dendritic integration. Baseline spike rates could not be recovered by lowering the dendritic h-channel density value alone to the value used in the younger model, and even decreased the spike rates further (**Fig. 3F**, paired-sample t-test, *p* < 0.001; Cohen’s *d*: -18.6). Thus, the age-associated decrease in spike rate was not due to changes in h-channels alone, but a combination of changes in both the depolarizing h-current and the hyperpolarizing leak current. Baseline spike rates could also not be recovered by changing the axo-somatic sodium, potassium, and calcium channel parameters in the older model to the values used in the younger model, which rather decreased the spike rates further (paired-sample t-test, *p* < 0.001; Cohen’s *d*: -13.2).

In line with decreased spike rates, the microcircuit models with older Pyr neurons also generated rhythmic population spiking with decreased peak frequency and power compared to the model with younger Pyr neurons (**Fig. 3G**). The spiking in both types of microcircuit models was in the beta frequency range, consistent with beta range frequency oscillations reported previously in local field potential recordings of L5 (Roopun et al., 2006).

To validate our model prediction that baseline spike rates of older human L5 pyramidal neuron microcircuits are reduced, we analyzed multi-electrode array (MEA) extracellular recordings across all layers of human cortical slices (**Fig. 4A-B**). These slices exhibited lower firing rates (**Fig. 4C**) compared to the simulated microcircuits (which modeled *in vivo* activity), but they exhibited the same dampening in older vs younger microcircuits. In the L5 areas of the slice, spike rates of broad-spiking single units (putative Pyr neurons) in older subjects were decreased compared to younger subjects (two-sample Welch’s t-test, *p* < 0.05; younger: mean = 1.09 Hz, 95% CI = 0.87 - 1.30 Hz; older: mean = 0.49 Hz, 95% CI = 0.30 - 0.64 Hz; Cohen’s *d*: - 0.27, respectively; n = 401 units from 14 younger subjects, n = 51 units from 4 older subjects; **Fig. 4C, top**). Similarly, spike rates in L2/3 areas of the slice were also decreased in older subjects compared to younger subjects (two-sample Welch’s t-test, *p* < 0.05; younger: mean = 1.27 Hz, 95% CI = 0.97 - 1.52 Hz; older: mean = 0.46 Hz, 95% CI = 0.31 - 0.57 Hz; Cohen’s *d*: - 0.37; n = 376 units from 14 younger subjects, n = 148 units from 5 older subjects; **Fig. 4C, bottom**). L5 rates were also found to decrease with age (Pearson correlation, R = -0.22, *p* < 0.05), but not L2/3 rates (**Fig. 4D**). Spike rates did not change significantly in L4 broad-spiking units with age, though this may be partly due to the presence of stellate neurons in L4. Compared to the 18.3% decrease in spike rates in our models with older L5 Pyr neurons due to sag amplitude changes alone, the L5 MEA data from older subjects showed a larger 55% decrease, but with a smaller effect size (Cohen’s *d*: -8.7 vs -0.27, respectively) due to the larger variability in the data. When averaged for each subject, neither L2/3 nor L5 rates decreased significantly with age, possibly due to the small subject sample size. We note, however, that linear mixed-effect models supported that the effects on spike rate were driven by age and not subject variability for both L5 and L2/3 (likelihood ratio test, *p* < 0.05).

**Figure 4.**
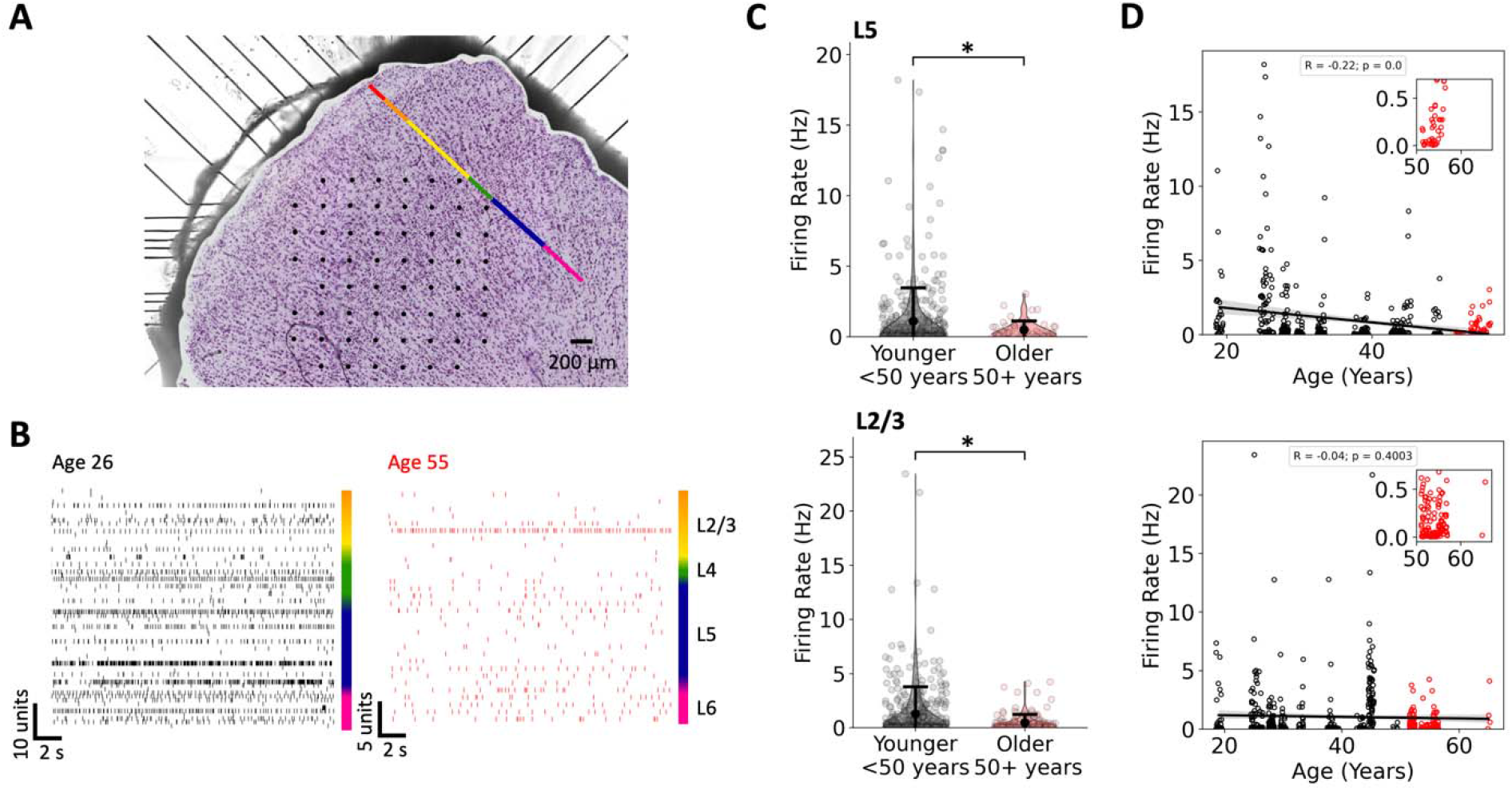
MEA recordings reveal dampened microcircuit spiking in older human cortical L5 and L2/3 Pyr neurons. **A**. Human cortical tissue (500 μm thickness; 5X lens) in a 60-channel MEA. Black dots denote electrode locations and the color bar denotes layers 1 (red, upper left) through 6 (pink, lower right) of the human cortical slice which was Nissl stained after the recording and overlaid on the image of slice during the recording. **B**. Example spike trains of single units ordered by cortical depth and grouped by layer (colour bar) from a younger subject (age 26, n = 65 units) and an older subject (age 55, n = 32 units). Units that were not spiking in this time range are omitted. **C**. Distribution of firing rates obtained from extracellular recordings by planar MEAs across layers of cortical slices. Error bars denote the bootstrapped mean firing rates and 95% confidence intervals. Firing rate was significantly decreased in older L5 (top) and L2/3 (bottom) Pyr neurons (two-sample Welch’s t-test). **D**. Firing rate measured in L5 (top) and L2/3 (bottom) neurons as a function of subject age. Inset plots show expanded axes for the older firing rate data. L5 neuron firing rates decreased significantly with age (Pearson correlation). The shaded area denotes the bootstrapped 95% confidence intervals.

### Increased sag amplitude has minimal effect on microcircuit stimulus response

To assess the effect of sag amplitude changes on microcircuit response, we examined tuning selectivity as a typical and generalizable example of cortical processing. We simulated orientation-selectivity in our models by applying thalamic inputs to orientation-selective neurons, with tuning curves as reported experimentally from monkey somatosensory cortex (**Fig. 5A, B**). We simulated different presentations of 85° or 95°, representing a small orientation difference, and thus a challenging discrimination task due to similar levels of circuit activation (**Fig. 5B**). Stimulus inputs included randomized additive noise and were applied to a microcircuit with fixed connectivity but random background input (**Fig. 5C-E**). Across 80 stimulus presentations for each angle, response rates of Pyr neurons were reduced in microcircuits with older Pyr neurons (paired-sample t-test, *p* < 0.05; younger: 10.39 ± 0.66 Hz; older: 10.19 ± 0.64 Hz; Cohen’s *d*: -0.3, **Fig. 5F**), although to a much lesser degree than the decrease in baseline rates (paired-sample t-test, *p* < 0.001; younger: 3.25 ± 0.03 Hz; older: 2.64 ± 0.03 Hz; Cohen’s *d*: -19.8). The larger dampening of baseline activity compared to response thus imposed an increased signal-to-noise ratio (SNR) in microcircuits with older Pyr neurons compared to those with younger Pyr neurons (**Fig. 5G**; paired-sample t-test, *p* < 0.001; younger: 3.06 ± 0.22; older: 3.68 ± 0.27; Cohen’s *d*: 2.5). Pyr neuron sag amplitude changes thus did not account for reduced response rates and decreased SNR seen in aging, indicating the involvement of other mechanisms.

**Figure 5.**
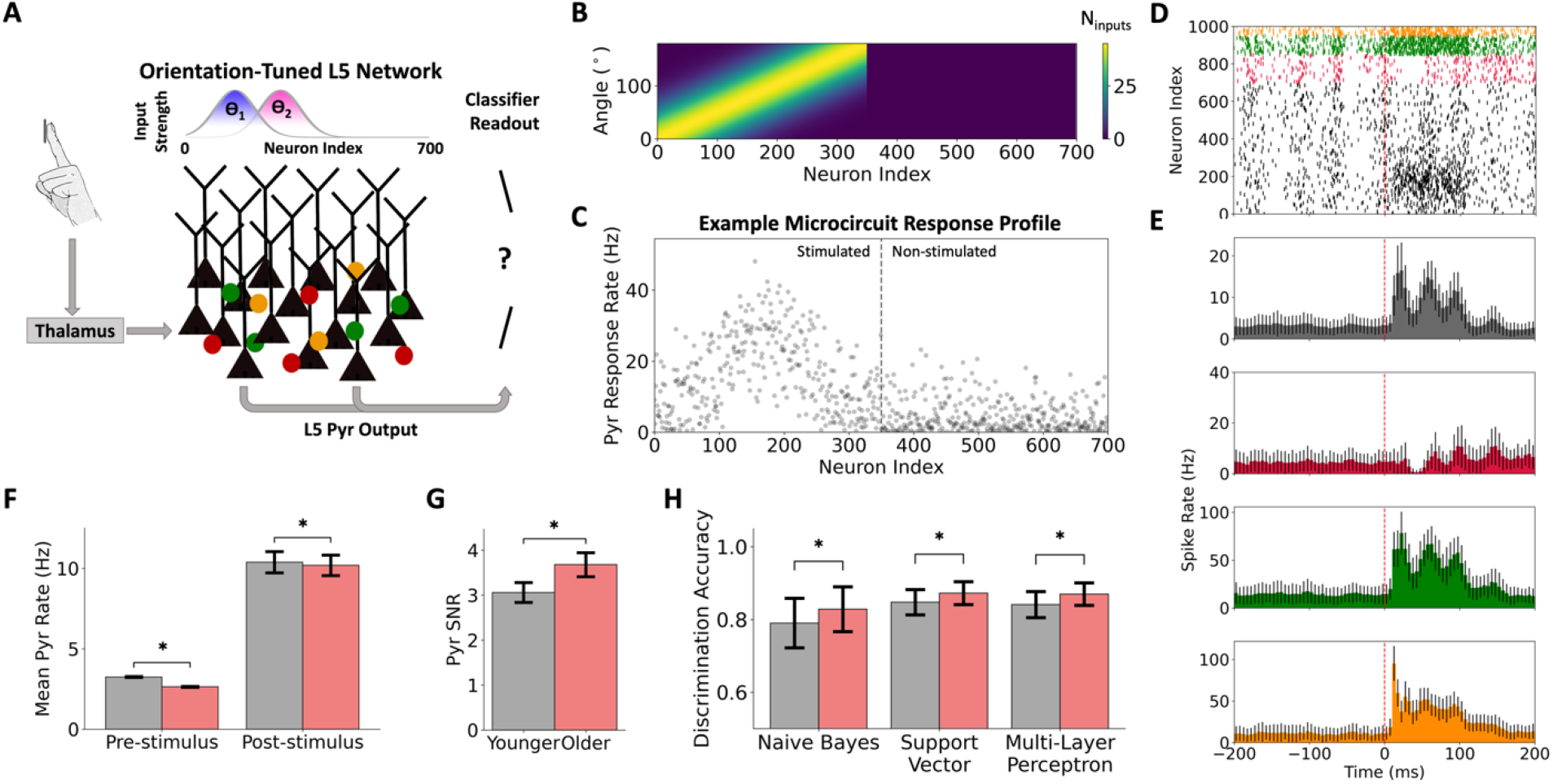
Increased sag amplitude in older Pyr neurons has minimal effects on stimulus response. **A**. Stimulus response simulation paradigm, where orientation-selective neurons in the L5 microcircuit received thalamic inputs corresponding to tactile edges of different orientations (e.g. □_1_, □_2_). The Gaussian distributions show the activation of different subgroups of neurons in response to different orientations. L5 Pyr neuron output responses to randomized stimulus presentations for each stimulus orientation (85° or 95°) were inputted into classifiers to assess discrimination accuracy. **B**. Orientation-tuning of Pyr neurons in the microcircuit, relating the number of thalamic inputs to each tactile edge orientation. **C**. Example average response rate for neurons in a microcircuit with younger Pyr neurons during (n = 80) presentations of the 85° stimulus. **D**. Example raster plot of simulated spike response in a microcircuit model with younger Pyr neurons during presentation of an 85° stimulus. Dashed line denotes the stimulus time. **E**. Average peri-stimulus time histograms during (n = 80) 85° stimulus presentations in the microcircuit model with younger Pyr neurons. **F**. Baseline and response spike rates to 85° inputs (mean ± SD) in both microcircuit models (n = 80 stimulus presentations each). **G**. Moderately increased SNR in cortical microcircuit models with older Pyr neurons compared to those with younger Pyr neurons. **H**. Moderately increased stimulus classification accuracy using readout from cortical microcircuits with older Pyr neurons compared to those with younger Pyr neurons, across different classifier types (n = 300 permutations of the train/test datasets).

We next determined the implications of SNR on the accuracy of discrimination of stimulus orientations using readout from microcircuits with older vs. younger Pyr neurons. The readout comprised of Pyr neuron output metrics that served as feature inputs into different classifier types (**Fig. 5H**). Across all classifiers, we found a moderate increase in discrimination accuracy when using output features from microcircuit models with older Pyr neurons compared to the microcircuit models with younger Pyr neurons (**Fig. 5H**; two-sample t-test, *p* < 0.05; Cohen’s *d* values: Naïve Bayes = 0.6; Support Vector = 0.7; Multi-Layer Perceptron = 0.9). The increase in microcircuit SNR mediated by the changes in sag amplitude mechanisms in older L5 Pyr neurons therefore led to some improvement in decoding signals from the neuronal activity readout. We also ran similar tests using spike rate or maximum PSP amplitude as alternative feature inputs, and these generated results that were consistent with our results when using area under the curve. We next examined Pyr neuron responses to apical dendrite inputs, as well as lower stimulus magnitudes (**Fig. 6A**). In all cases, changes to response rates were minimal (**Fig. 6B**; Cohen’s *d*, paired-sample t-test: low basal = -0.8, *p* < 0.01; low apical = -1.3, *p* < 0.001; basal = -0.6, *p* < 0.01; apical = 0.3, *p* > 0.05) and SNR was higher in microcircuit models with older Pyr neurons (**Fig. 6C**; paired-sample t-test, *p* < 0.001, Cohen’s *d*: low basal = 1.6; low apical = 1.4; basal = 2.0; apical = 2.9). These effects on SNR were smaller for responses to weaker inputs (**Fig. 6C**) due to larger reductions in response rates with age in this condition (**Fig. 6B**). Thus, in all simulated tests, Pyr neuron sag amplitude changes alone could not sufficiently account for the reductions in response rates and discrimination acuity seen experimentally in aging.

**Figure 6.**
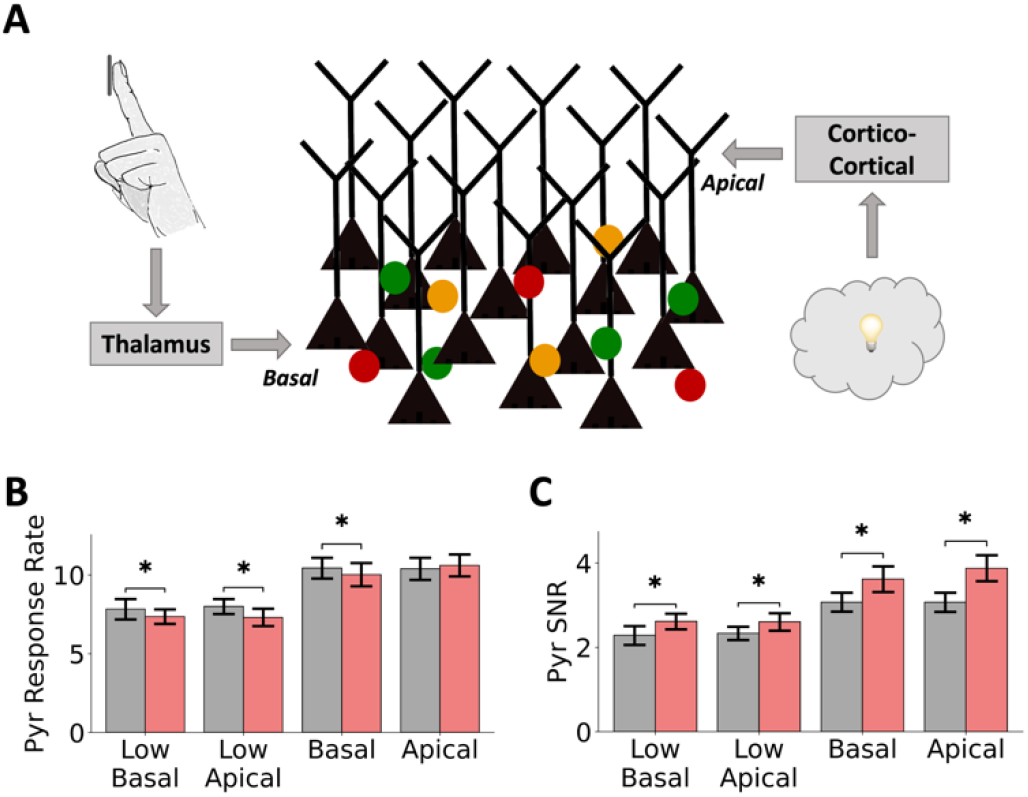
Age-associated changes are robust across stimulus location and amplitude. **A**. Stimulus response simulation paradigm, where the Pyr cells are stimulated by either thalamic inputs (representing tactile edges) onto their basal dendrites, or cortical inputs (representing associative information) onto their apical dendrites. **B**. Minimal effects on response rates in cortical microcircuit models with older Pyr neurons compared to those with younger Pyr neurons (n = 30 stimulus presentations) for low/high input onto basal/apical dendrites. **C**. Increased Pyr neuron SNR in cortical microcircuit models with older Pyr neurons compared to those with younger Pyr neurons.

As the differences in excitability between younger and older Pyr neuron models were primarily driven by changes in dendritic h-channel density and membrane leak current parameters (**Fig. 3E**), we examined excitatory (**Fig. 7A**) and inhibitory (**Fig. 7D,G**) PSP summation in the models to characterize possible reasons for the dampened firing rates in microcircuits with older Pyr neurons. The peak excitatory (**Fig. 7B, top**) and inhibitory (**Fig. 7E,H, top**) PSPs (from Pyr and SST/PV connections, respectively) in the older neuron model were more depolarized than in the younger neuron model, due to the depolarized resting membrane potential resulting from larger h-channel density. In addition, the inhibitory PSP amplitude of the PV→Pyr connection increased considerably (paired-sample t-test, *p* < 0.05, Cohen’s *d*: 5.4; **Fig. 7I, left**). While we also observed decreased Pyr→Pyr (paired-sample t-test, *p* < 0.05, Cohen’s *d*: -0.6) and increased SST→Pyr (paired-sample t-test, *p* < 0.05, Cohen’s *d*: 0.2) amplitudes (**Fig. 7C,F, left**), these effects were small. When current-compensated to equalize resting membrane potential between the models, excitatory (**Fig. 7B, bottom**) and inhibitory (**Fig. 7E,H, bottom**) PSP summation in the older neuron model were dampened compared to the younger neuron model (paired-sample t-test, *p* < 0.05, Cohen’s *d*: Pyr→Pyr = - 1.9, SST→Pyr = -1.7, PV→Pyr = -1.5; **Fig. 7C**,**F**,**I, right**). Across all connection types, the PSP time course was shortened in the older neuron model, reflecting the larger h-current effects. Although the net effect on IPSP and EPSP summation in the microcircuit depends on the circuit state and neuronal membrane potential, these results indicate dampened excitation and boosted inhibition in older Pyr neurons which would generally account for the reduced firing rates. We obtained similar significant effects also when using a lower stimulation frequency of 25 Hz (paired-sample t-test, *p* < 0.05, Cohen’s *d*: Pyr→Pyr = -0.5, SST→Pyr = 0.25, PV→Pyr = 4.8), as well as during 25 Hz stimulation with current-compensation to equalize resting potentials (paired-sample t-test, *p* < 0.05, Cohen’s *d*: Pyr→Pyr = -1.3, SST→Pyr = -1.3, PV→Pyr = -1.3).

**Figure 7.**
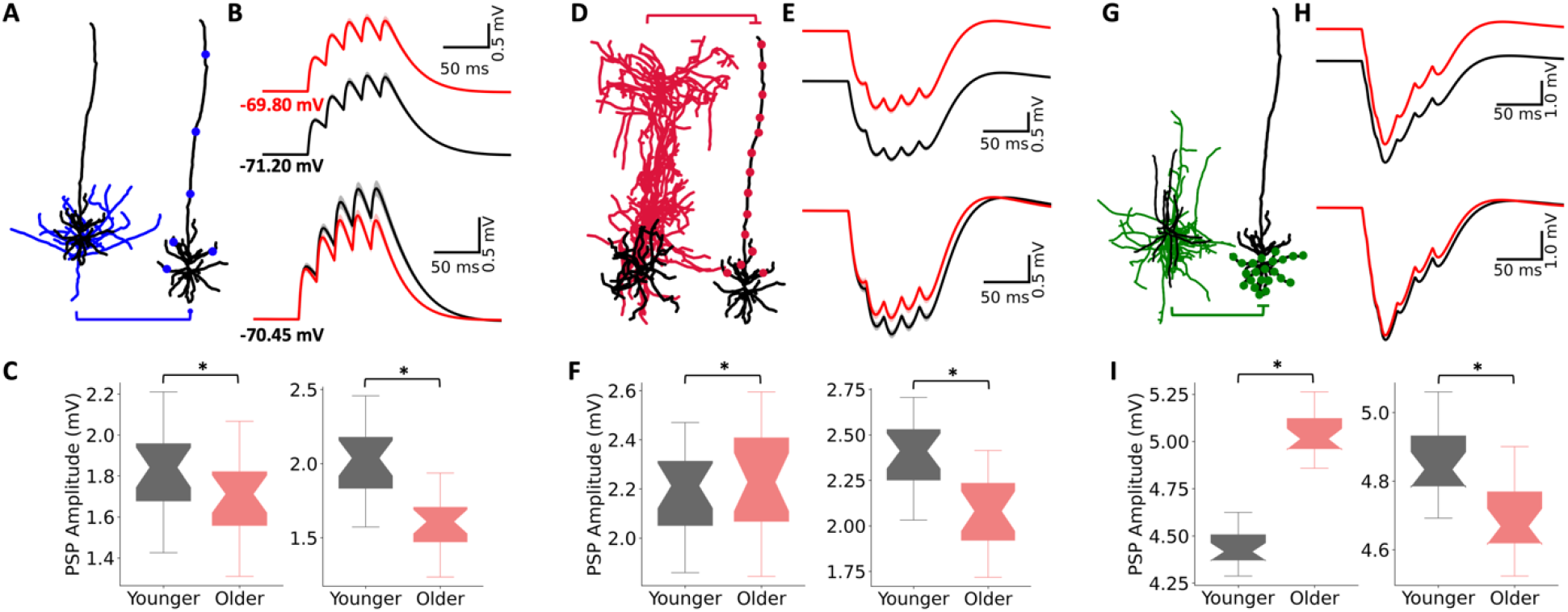
Synaptic input summation in younger and older neuron models. **A**. Pyr→Pyr connections tested for EPSP summation. Circles mark synaptic contacts onto the post-synaptic Pyr neuron. **B**. Top traces show EPSP summation (50 Hz train of 5 step current pulses) in the younger (black) and older (red) neuron models (20 connections, 20 trials/connection), bootstrapped mean and 95% confidence intervals. Bottom traces show the same as the top traces, but when resting membrane potential is equalized between younger and older neuron models by current compensation. **C**. Summary statistics of peak PSP amplitude in younger and older Pyr neuron models corresponding to the traces in B,top (left) and B,bottom (right). The box boundaries denote the interquartile ranges, the notches in the boxes denote the medians, and the whiskers denote the ranges. **D-I**. Same as A-C but for IPSP summation of SST→Pyr (D-F) and PV→Pyr (G-I) connection types.

## Discussion

Here we found an increase in sag amplitude of human cortical L5 pyramidal neurons with age, which led to reduced baseline activity and noise during response in simulated human cortical microcircuits. Analysis of extracellular recordings using MEA across layers *ex vivo* further confirmed a decrease in L5 pyramidal neuron spike rates in cortical tissue from older individuals. We demonstrated the link mechanistically using novel detailed models of human L5 microcircuits that integrated human cellular and circuit data. Our simulations show that although changes in Pyr neuron sag amplitude mechanisms did not play a role in age-dependent reduced response rates and discrimination acuity, the sag amplitude changes alone can account for the reduced baseline spike rates seen in aging. Our models thus reproduce age-dependent changes in cortical resting state activity, and may serve to further study their implications and clinical relevance in aging.

The reduced baseline and response rates in our microcircuit models in aging, and the involvement of h-currents, agree with previous studies in monkeys (Wang et al., 2007, 2011). Two key differences in the above studies, however, were a larger reduction in response rates accompanied with cognitive deficits in working memory, and a global suppression of h-channels rescuing both response spike rates and working memory deficits. The further reductions in spike rates that we observed when simulating reduced Pyr neuron h-channel conductance and the improved discrimination accuracy with older Pyr neurons were more in line with the effects of HCN1 deletion or blocking h-current that have been reported for mouse prefrontal cortex (Thuault et al., 2013). While the effects of h-current suppression on prefrontal cortex spiking and working memory remain to be clarified, our findings thus indicate that the effects of global h-channel suppression on circuit spiking in monkeys likely go beyond the impact in Pyr neurons that we have modelled here. These discrepancies may be due to multi-regional effects, or the involvement of age-changes in other mechanisms as indicated by our simulations of response and discrimination acuity. Possible additional age-associated mechanisms through which reduction in response rates and cortical processing quality could occur include reduced SST and VIP inhibition (Mohan et al., 2018), neuronal atrophy (Prevot et al., 2021), homeostatic reductions in tonic inhibition (Chen et al., 2010; Prevot et al., 2021), and reduced NMDA/AMPA ratios (Pegasiou et al., 2020). There may also be sag amplitude changes in other neuron types such as SST interneurons, which exhibit large sag amplitude like Pyr neurons, but for which aging data is not yet available. The possible involvement of the above mechanisms may also depend on associative distal dendritic inputs onto L5 pyramidal neurons, which we demonstrated can exhibit similar effects on baseline and response rates due to increased sag-current in the microcircuit models with older Pyr neurons. In this work, although we focused on group effects in line with previous studies (Bucur & Madden, 2010; Davis et al., 1990; McIntosh et al., 2014), we also reported significant correlations between both L5 Pyr neuron sag amplitudes and spike rates with age. In both comparisons, the age effect on sag was stronger when examined at larger hyperpolarizing current steps or across current steps. Further studies and protocols would benefit from ensuring a large range of hyperpolarizing step amplitudes. Our comparison across neurons rather than subjects is supported by the similar inter- and intra-subject neuronal data variability, and by the effects not being driven by any one individual, but it will also be of interest in future studies to increase the subject sample size and thus better establish the effect across subjects rather than neuron samples. Analysis of *ex vivo* MEA data also validated our predictions of decreased L5 pyramidal neuron spiking in cortical microcircuits of older individuals. While the decrease in microcircuit spiking was greater than the changes seen in our microcircuit models involving sag amplitude change alone, differences in the magnitude of the decrease could be attributed to the more silent state of the *in vitro* tissue (which is therefore more susceptible to dampening mechanisms) and/or the involvement of other mechanisms mentioned above. Whereas in the current study we have assessed the functional impact of Pyr neuron changes with age, it will be of interest in future studies to explore how additional diverse mechanisms refine our results to explain the rest of the effects seen in aging.

We studied the effect of the age-dependent cellular changes on sensory cortical processing, since the underlying microcircuits are well-studied (Yu et al., 2019; Thakur et al., 2006; Bensmaia et al., 2008) and since sensory processing worsens with age. Previous studies showed that tactile acuity decreases with age (Legge et al., 2008, 2019), and perceptual speeds and reaction times are slower (Bucur & Madden, 2010). Though we constrained the *in vivo* aspects of our model using rodent and monkey data due to unavailability of corresponding human data (**Table S6**), the backbone of our models (i.e., morphologies, neuronal excitability, cell type proportions, Pyr to Pyr connectivity) was constrained by human data and so we expect our findings to be relevant to the neurophysiology of human circuits. Furthermore, the simulation of cortical response to sensory stimuli served to study a prototypical cortical microcircuit processing to bottom-up inputs together with recurrent inputs from the microcircuit neurons onto Pyr neuron basal and apical dendrites (Hay & Segev, 2015). The consistent age effects we saw when simulating the processing of apical inputs, which would correspond to cortico-cortical connections from higher-order cortical areas, indicate that the results should generalize to other types of cortical processing relevant to aging.

The age-dependent reduction in microcircuit spike rates involved a combination of dendritic mechanisms underlying the increased sag amplitude, including h- and leak-currents. Despite the older models being more depolarized, circuit spiking was reduced due to increased h-current dampening synaptic summation, in agreement with previous studies (Williams & Stuart, 2000; Song & Moyer, 2017), and due to a reduced excitatory driving force and an increased inhibitory driving force. While we modeled the altered sag amplitude via a change in h-channel density proportionally along the apical dendrites, future experiments should determine if there is also a change in the dendritic distribution of h-channels, which may impact the proportion of proximal vs distal densities and thus differentially affect input integration. The increase in h-current estimated from our models and electrophysiology was further supported by analysis of HCN1 expression in human Pyr neurons. HCN channel subunits are expressed ubiquitously across layers of human cortex, more than in rodents, further suggesting their importance in human Pyr neurons (Kalmbach et al., 2018). While most of the data we used was healthy tissue obtained from people with epilepsy (Chameh et al., 2021), HCN channels can be up- or down-regulated in epilepsy depending on the brain area and/or species (Albertson et al., 2011; Arnold et al., 2019; Bender et al., 2003), and we cannot discount the possible influence of this in our findings. A notable difference between the KBI and ABI datasets was the use the synaptic blockers in the ABI dataset, which can account for the larger (∼2-fold) input resistance values in that dataset. Though our findings were mostly conserved across the two datasets, future studies can investigate the effects of synaptic currents which can enhance the recruitment of dendritic h-currents (Guet-McCreight & Skinner, 2020). Although we have studied the effect of Pyr neuron sag amplitude, it remains unknown whether there are also age-associated changes in interneuron sag amplitude, which would modulate cortical firing rates. SST, PV, and VIP -expressing interneurons have all demonstrated h-currents in electrophysiological recordings (Albertson et al., 2017; Prönneke et al., 2015; Roth & Hu, 2020). When human interneuron electrophysiology data becomes more abundant, a similar approach could be used to estimate age-associated changes and integrate them into the models to better understand the microcircuit changes in aging. We also note that action potential width in our models was narrower than the experimental ranges due to the limitations imposed by the set of ion channel mechanisms used in this study and previously (Druckmann et al., 2007; Hay & Segev, 2015; Mäki-Marttunen et al., 2019; Markram et al., 2015). Thus, though these models were sufficient in capturing neuronal spiking patterns and input-output gain for network simulations, future improvements to the models could expand sodium and potassium channel kinetics to better capture spike width features.

Our microcircuit models capture several aspects of biological L5 circuits, including human cell type proportions, human neuronal intrinsic properties, *in vivo* spike rates, and stimulus response rates. The Pyr neuron models in our study were constrained with both morphological and spiking features of intratelencephalic-projecting-like neurons, in line with the higher proportion of these neurons in human MTG relative to extratelencephalic-projecting Pyr neurons (Chameh et al., 2021; Hodge et al., 2019; Kalmbach et al., 2021). We did not have data for the *in vivo* spike rates of different L5 Pyr neuron subtypes (e.g. intratelencephalic-projecting type 1 and type 2, and extratelencephalic-projecting) and the proportions of these neuron types vary across species (Hodge et al., 2019; Kalmbach et al., 2021), therefore we treated L5 Pyr neurons as one population in our models. However we note our model morphologies are intratelencephalic-projecting-like, and the data we fit our models to is likely predominantly recorded from intratelencephalic-projecting Pyr neurons based on their electrophysiology metric values (**Figs. S2 and S3**; Kalmbach et al., 2021). When dendritic properties of human L5 intratelencephalic-projecting Pyr neurons become better understood (for examples of works that include dendritic characterization in other types of human Pyr neurons see: Beaulieu-Laroche et al., 2018, 2021; Gidon et al., 2020; Kalmbach et al., 2021), it will be interesting to investigate the possible contribution of intratelencephalic-projecting L5 Pyr neuron dendritic features such as action potential backpropagation and calcium spikes to age changes in dendritic h-currents. For example, these could amplify the effect on network response rates, a feature that has been demonstrated with rodent L5 extratelencephalic-projecting Pyr neuron network models (Hay & Segev, 2015). Whereas we simulated L5 microcircuits of a smaller size than the true size (by a factor of ∼3), which decreases the overall recurrent inputs received by each neuron in the network, we confirmed with simulations of larger microcircuits that spiking activity was similar and our results held. This is due to our microcircuit down-sampling involving proportional increases in both excitatory and inhibitory neurons, which maintained the overall excitatory-inhibitory balance of the network. Our circuit models spiking exhibited beta range frequency oscillations, which have been reported in local field potential recordings of L5 (Roopun et al., 2006; Halgren et al., 2018). The origin of this rhythm, which was not a feature that we explicitly tuned our models to exhibit, could be the product of many components, but we expect is primarily governed by the recurrent excitatory and inhibitory connections in the circuit, since the background input is random. Also, the larger proportion of PV interneurons (e.g. compared to L2/3) could further explain the oscillations at these particular frequencies (Cardin et al., 2009). The slowing of the peak frequency and reduced power with age in our models may also bear similar mechanisms to the slowing and decreased power of alpha rhythms with age that have been reported in EEG datasets (Choi et al., 2019; Donoghue et al., 2020). Future models involving also the superficial cortical layers would enable linking the age-dependent changes to the associated EEG signals (Hagen et al., 2018), and relating them to experimental recordings (Choi et al., 2019; Donoghue et al., 2020). All models and simulation code will be available openly online upon publication.

## Acknowledgements

AGM, SJT and EH thank the Krembil Foundation for their generous funding support. MW and EH thank NSERC for funding support. SJT and TAV also thank the generous support from CAMH Discovery Fund and Kavli Foundation. We also thank Dr. Yuxiao Chen for technical assistance in gene-expression analysis. As well, we are immensely grateful to our neurosurgical patients and their families for consenting to the use of their tissue samples for research.

